# Phase separation as higher-order catalyst

**DOI:** 10.1101/2022.10.27.514140

**Authors:** Kai Huang, Xuebo Quan, Shiyi Qin

**Author notes:** Correspondence (K.H.).

## Abstract

The long-distance communication between multiple cis-regulatory elements (CREs), the self-limiting size and lifetime of regulatory condensates, are two puzzling phenomena in biology. To reconcile these puzzles, we introduce the concept of higher-order catalysis into chromatin-mediated reactions. Essentially, multi-way contact between the CREs defines a transition state that is required for the downstream cascade of chemical reactions. The entropic penalty of chromatin reorganization sets a high activation barrier to enter this transition state. Phase separation of trans-acting agents induced by the CREs reduces this barrier and stabilizes the transition state via forming a regulatory condensate. The downstream reaction then pays back energy to dissolve the condensate and resets the agents to a metastable single-phase state. Accelerating the reactions without consuming agents or changing their state, the cycled phase transitions construct a higher-order catalyst or super-enzyme that is beyond the form of a single molecule. We discuss how chromatin employs such super-enzymes to catalyze higher-order reactions mediated by itself.

## Introduction

For unknown reasons, cis-regulatory elements (CREs) in mammalian cells often have admirable distance from their targets. It is a long-standing mystery how these elements tens or hundreds of kilobases away from each other communicate and coordinate their activities in space and time^1^. Cohesin-driven loop extrusion, an energy-consuming mechanism, can provide partial explanation to the contact formation between distal CREs^2,3^. However, perturbation or even eradication of loop extrusion seem to have limited effects on the global profile of gene expression^4,5^, indicating the existence of alternative mechanisms that can also contribute to CRE interactions for robust transcriptional regulation. It has been long known that the transcription enzyme RNA polymerase II (Pol II) prefers to aggregate in distinct hubs, a phenomenon that led to the proposal of the transcription factories model dated back to the 1990s^6^. Some recent studies indicate that these Pol II hubs can harbor multiple CREs and are liquid droplets likely formed by a mechanism called liquid-liquid phase separation (LLPS)^7–11^. An archetypal example of LLPS is the immiscible solution of water and oil. As an old concept in physical chemistry, LLPS is in its new dawn in biology, attracting growing research interest^12,13^. While a wealth of biomacromolecules including the CRE-binding transcription factors and cofactors have been reported to undergo LLPS in test tubes, their forms of organization inside live cells are highly complicated and therefore subject to controversy^14–18^. Deeper understanding is in urgent need to place LLPS, a thermodynamically spontaneous process, in the nonequilibrium context of cell biology, so that biophysical mechanisms can be better linked to functional roles.

The most widely recognized role of LLPS is to constitute a membrane-less organelle (MLO) usually hundreds to thousands nanometer large that enrich a selection of molecular species for specific biological functions such as biological synthesis^19^, stress response^20^, and signal transduction^21^. A typical nuclear MLO is the nucleolus, a macroscopic liquid condensate with clear function, i.e., to assemble ribosomes^19^. As a master of LLPS, nucleolus uses multiple liquid phases to build its assembly line that allows centralized and efficient ribosome biogenesis, which makes itself a paradigmatic example of a transcription factory. However, when it comes to the Pol II enriched CRE hubs, we confront a fundamentally different scenario with puzzling observations that challenge our current understanding of LLPS and transcription. First, the CRE hubs are large in number, highly decentralized, and widely distributed in the nucleus. On the length scale, most of them are 100 nanometer or less in diameter, significantly smaller than typical MLOs^22,23^. Such abnormal size fuels the debate on the formation mechanism of the CRE hubs^14^. If LLPS is the driving force, it is a mystery how the CRE hubs manage to escape droplet coalescence and Ostwald ripening^24^ to coexist with self-limiting sizes. Moreover, while LLPS is thermodynamically stable, many CRE hubs have lifetimes as short as just a few seconds^22,23^. This is the timescale of transcribing about only 1000 base pairs without even accounting for the transcription pausing that can take much longer time^25^. The unique spatiotemporal properties of the CRE hubs make them apparent outliers of the MLO paradigm and prompts us to rethink their biological function.

Zooming out from the scale of the CRE hubs to the 4D genome, we here seek to understand not only the mechanisms underpinning the assembly and disassembly of the dynamic structures, but also how their life cycle can be integrated into a higher-order system that regulates gene expression. Aware of the daunting complexity of the regulatory machinery, we leave out all the molecular details to focus on the very basic principles of physical chemistry that apply to both non-living and living systems, asking the question how evolution could have taken advantage of these rules for better control of genomic information. We first acknowledge that chromatin takes the form of a long fiber, whose conformations are heavily shaped by entropic forces, according to polymer physics^26^. A classic example of entropic force is the elasticity of freely jointed polymers. Despite its softness, a freely jointed polymer under a stretched condition tends to collapse to maximize its conformational entropy. In the same spirit, forming a loop, especially a large one, is entropically unfavorable and therefore unlikely to happen by chance. The possibility of long-range multi-way contacts that reorganizes chromatin into multi-loops is even lower if no external driving forces are present. While traditional Hi-C methods are limited to detection of pairwise chromatin contacts^27^, advances in experimental techniques allow us to see the existence of multi-way genomic interactions^7,28–31^. On the theoretical side, we have recently developed a three-dimensional (3D) forest model of chromatin^32,33^ to reconcile the paradox between long-range interaction^34^, packing heterogeneity^35^ and self-similar compartmentalization^36^, which counterintuitively predicts that multi-way contacts not only exist but prevail at the single-cell level. While our phenomenological model explains many structural features of chromatin, the molecular mechanisms of the predicted multi-way genomic interactions remain to be addressed. To answer this question, it is instructive to classify genomic contacts into functional ones and nonfunctional ones. Take the loop extrusion model^2,3^ for example, the active extrusion process keeps creating pairwise chromatin contacts in search for the functional ones. While this scanning strategy can boost the probability of forming functional contacts, most of the energy is consumed on creating contacts that have no functions. Given the fact that enhancers greatly outnumber promoters^1^, functional multi-way contacts involving several enhancers might not be less common than one-on-one enhancer-promoter pairs. However, as the number of possible *N*-body contacts increases rapidly with *N,* it would be impractical for the chromatin machinery to scan all the multi-way interactions. Recalling the high entropic penalty associated with multi-way long-range interactions on polymers, molecular mechanisms that selectively and efficiently establish functional multi-way genomic contacts would be desirable from an energy economics point of view.

In this work, we propose a higher-order catalysis model to shed light on the mechanism and function of multi-way genomic interactions. We show that LLPS of trans-acting agents is an efficacious way to drive the formation of CRE hubs. To make sense of these short-lived tiny condensates, we reason that they are not playing the traditional role of stable MLO for transcription-related reactions, but rather a stabilized transition state of a higher-order reaction for gene regulation. We propose that the conformational entropy of chromatin polymer and the enthalpy-driven LLPS of agents are two antagonizing forces that can work in synergy to construct a higher-order catalyzing system in which the entropic force sets up the activation barrier that limits the reaction rate whereas the enthalpic force selectively lowers the barrier to accelerate the higher-order reaction. Once the cascade of downstream genomic reactions is triggered, part of the released energy can be used to dissolve the condensate so that the agents transition back to a metastable single phase. We employ a minimal computational model to quantitatively study the clustering of CREs driven by LLPS and compare it with a bridging mechanism that we find to be less effective in mediating multi-way interactions. We discuss the implication of our theoretical model and the molecular insights from our simulations.

## The higher-order catalysis model

Gene regulation involves a plethora of chemical reactions mediated by chromatin. In our model, we coarse-grain a series of chemical reactions and related physical processes into a higher-order regulatory reaction. A typical higher-order reaction of our interest has a few CREs as its reactants and gene expression or silencing as its products. The associated physical processes include diffusion and binding of trans-acting molecules, the clustering of the CREs and the concomitant looping of chromatin. Taking advantage of the timescale separation, we here only concern the slow dynamics of chromatin, neglecting the fast motion of small molecules (compared to chromatin) and their binding kinetics. We do not explicitly account for the conformational freedom of chromatin, but rather coarse-grain it into the free energy landscape of the higher-order reaction. Once the chromatin arranges itself so that the CREs as reactants cluster in space, the higher-order reaction can proceed with a certain rate. However, based on polymer physics, such reorganization of chromatin is associated with a penalty of free energy Δ*G* as the CRE hub reduces the conformational freedom of chromatin. Therefore, in the language of transition state theory^37^, the clustering of CREs defines a transition state or activation complex that limits the rate of the reaction. The overall higher-order reaction can be written as:

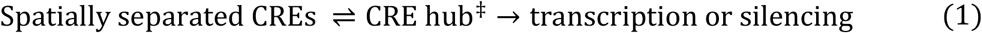

where the symbol ‡ denotes the transition state. Here for simplicity, we ignore non-CRE elements such as transcription factors, cofactors, Pol II, chromatin remodelers, noncoding RNAs and ATPs. The activation rate constant of the higher-order reaction *K*_A_ depends on the activation free energy Δ*G* in an exponential way:

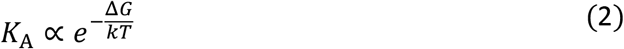

Here, *k* is the Boltzmann constant and *T* is the temperature. Note we use the Boltzmann constant instead of the universal gas constant because the higher-order reactions in single cells are individual events with a small number of reactants and products.

Now that we have developed the framework of higher-order genomic reactions and generalized the concepts of transition state and activation barrier, it is natural to ask what mechanism can act like a catalyst to increase the reaction rate. We show that LLPS, the thermodynamic force that builds macroscopic MLOs, can be used to make a higher-order catalyst. It is well known that phase transition does not necessarily happen beyond the critical point if no nucleation sites are provided^38^. Supercooled water is a classic example as it stays in liquid state below the freezing point in the absence of seed crystal. Likewise, nucleation is needed for proteins to undergo LLPS when the protein concentration falls in between the binodal (coexistence) and spinodal^39^ curves. In this regime, the system can be kinetically trapped in a metastable single-phase state even though the two-phase state is thermodynamically more favorable. The metastability allows the phase behavior to be responsive to nucleation stimuli, which makes sensitive higher-order catalyzing possible. In the higher-order genomic reaction, the nucleation stimuli can be the transcripts of CRE-associated non-coding RNA (ncRNA) such as enhancer RNA (eRNA)^40^. It has been uncovered that a great portion of nuclear proteins can bind to ncRNAs^41^. A large pool of ncRNAs and their cognate nuclear proteins could endow good catalyst selectivity. Once the nucleation capacity of CREs is turned on, LLPS of selected transacting agents will be induced. When the enthalpic gain of the LLPS is large enough to overcome both the cost of translational entropy of the agents and the penalty of conformational entropy of chromatin, the condensation will pull together the nucleation sites, leading to the assembly of the CRE hub. As such, the induced LLPS stabilizes the activation complex of the higher-order genomic reaction. We use the term of regulatory condensate instead of transcriptional condensate to describe the dense phase since in our model the condensate regulates the rate of the higher-order reaction rather than making the end products such as messenger RNA (mRNA). Regarding catalyst efficiency, we propose that the higher-order reaction will not pause at the transition state for too long before the downstream chemical reactions dissolve the regulatory condensate to regenerate the single-phase state that is ready for the next catalytic action. It is worth noting that since the phase-separated state is thermodynamically more stable than the single-phase state, the induced LLPS cannot be reversed by simply turning off the nucleation sites. The dissolution of the regulatory condensate entails a gain of energy which can be supplied by chemical reactions such as the degradation of the ncRNAs and the post-translational modification (PTM) of Pol II. Ideally, the amount of energy needed for condensate dissolution is equal to the mechanical work done by the LLPS to cluster up the CREs. As schematically shown in Figure 1, the collective action of the agents during the cycle of phase transitions speeds up the higher-order reaction without changing the free energy difference between the reactants and products. This is a typical catalytic process since no agents are consumed and their state remains the same after the higher-order reaction. Given its supramolecular nature and the large scale of its substrate, we use the term higher-order catalyst or super-enzyme to describe the accelerating machinery.

**Figure 1.**
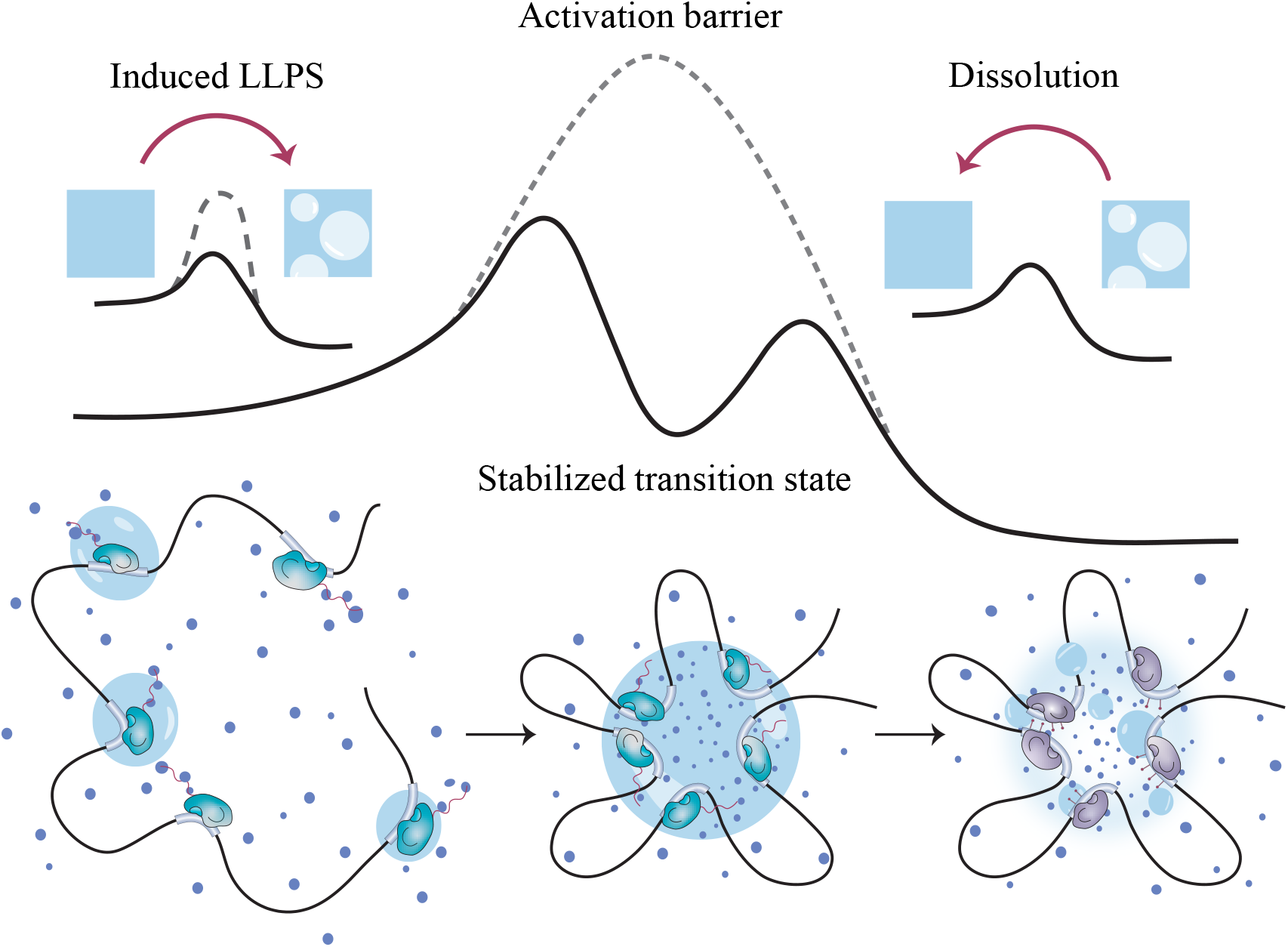
Schematic illustration of the higher-order catalysis model. Multi-CRE-regulated gene expression is used as an example of the higher-order reaction. We do not distinguish enhancers and promoters for simplicity. The separated CREs need to cluster up to activate the higher-order reaction. The free energy landscape of the higher-order reaction has a high activation barrier (dashed line) as it demands chromatin reorganization into multi-loops anchored by the CREs. LLPS of trans-acting agents induced by the CREs lowers this barrier (solid line) and brings the CREs into multi-way contact. The CRE hub associated with the regulatory condensate of the agents defines a stabilized transition state. The transition state then triggers downstream chemical reactions as part of the higher-order reaction to dissolve the regulatory condensate. The induction of the LLPS, the growth of the condensate to capture the CREs, and the dissolution of the droplet constitute an enzymatic cycle. The corresponding free energy landscapes of the agent phase transitions are shown above their corresponding schematic configurations at the different stages of the higher-order reaction. The dashed line represents the original high nucleation barrier without the induction of the CREs. Different colors of the Pol II in the schematic configurations highlight their change of chemical states during the higher-order reaction.

## Computer simulation results

Computer simulation is a powerful tool to investigate the dynamics of molecules, especially when the system is out of equilibrium. Since the activation barrier and the accelerating mechanism of the higher-order reaction mediated by chromatin can be well described by classical statistical mechanics, we choose Langevin dynamics^42^ as our simulation method for the sake of computational efficiency. We build a coarse-grained simulation system in which only part of the chromatin is considered, modeled as beads on a string confined in a cubic box filled with transacting agents. For simplicity, we consider only one kind of agent in our system which is modeled as a short chain of 5 monomers. Unless otherwise specified, 5 CREs, each 10-bead long, are dispersed on the chromatin fiber with the intervals being 1000 chromatin beads. We model inactive CREs as normal chromatin beads of weak cohesiveness and use relatively strong CRE-agent interaction to depict the active state of the CREs that can nucleate LLPS. All the bonds are modeled as harmonic and all the nonbonded interactions are described by Lennard-Jones potential. We first relax the system with inactive CREs before we turn on the specific CRE-agent interactions simultaneously. Under strong CRE-agent binding, agents can bridge multiple CREs together without LLPS. To investigate the difference between bridging and LLPS on mediating multi-way genomic contacts, we used two distinct combinations of CRE-agent and agent-agent interaction parameters. More simulation details can be found in the simulation methods section of the supporting information (SI). The minimal molecular model allows us to prove the concepts of higher-order catalysis as all the system variables can be well understood.

We first demonstrate that the entropic force of chromatin as a long polymer can lead to an energetic barrier for CREs to approach each other in space. For simplicity, we only consider two chromatin-connected CREs in an agent-free box and calculate the potential of mean force (PMF) as a function of their spatial distance. Figure 2A shows that the energetic barrier is long-range and scales with the length of the chromatin segment that separates the two CREs. Note that for ideal chains the height of the barrier can be analytically solved if one ignores the self-interaction of the polymer and the confinement effect. Our PMF calculations suggest that the barrier for two CREs with long sequence distance on chromatin to meet can be higher than 10kT. This is not a very strong barrier though, indicating that the pairing of two CREs alone can hardly limit the rate of regulatory reactions. When it comes to multi-way contact, as the barrier height for N-way contacts scales roughly with N-1, the cost of forming a CRE hub involving 5 or more CREs can be above 50kT, corresponding to a reaction time scale of days or even years. This is because the reaction rate constant *K*_A_ decreases exponentially as the activation free energy Δ*G* goes up (Equation 2). We then calculate the PMFs for two-body CRE interactions in the presence of agents that undergo induced LLPS and bridging, respectively. As shown in Figure 2B (and Figure S1, S2 with more details), LLPS leads to a long-range attraction with a shallow energetic well. The remaining barrier is weak for the CREs to enter the well. In stark contrast, bridging issues a very deep but narrow energetic well, with the remaining barrier being relatively larger. Although we did not assign strong attraction between the monomers of the CREs and the agents, the additive multivalent interaction leads to strong avidity.

**Figure 2.**
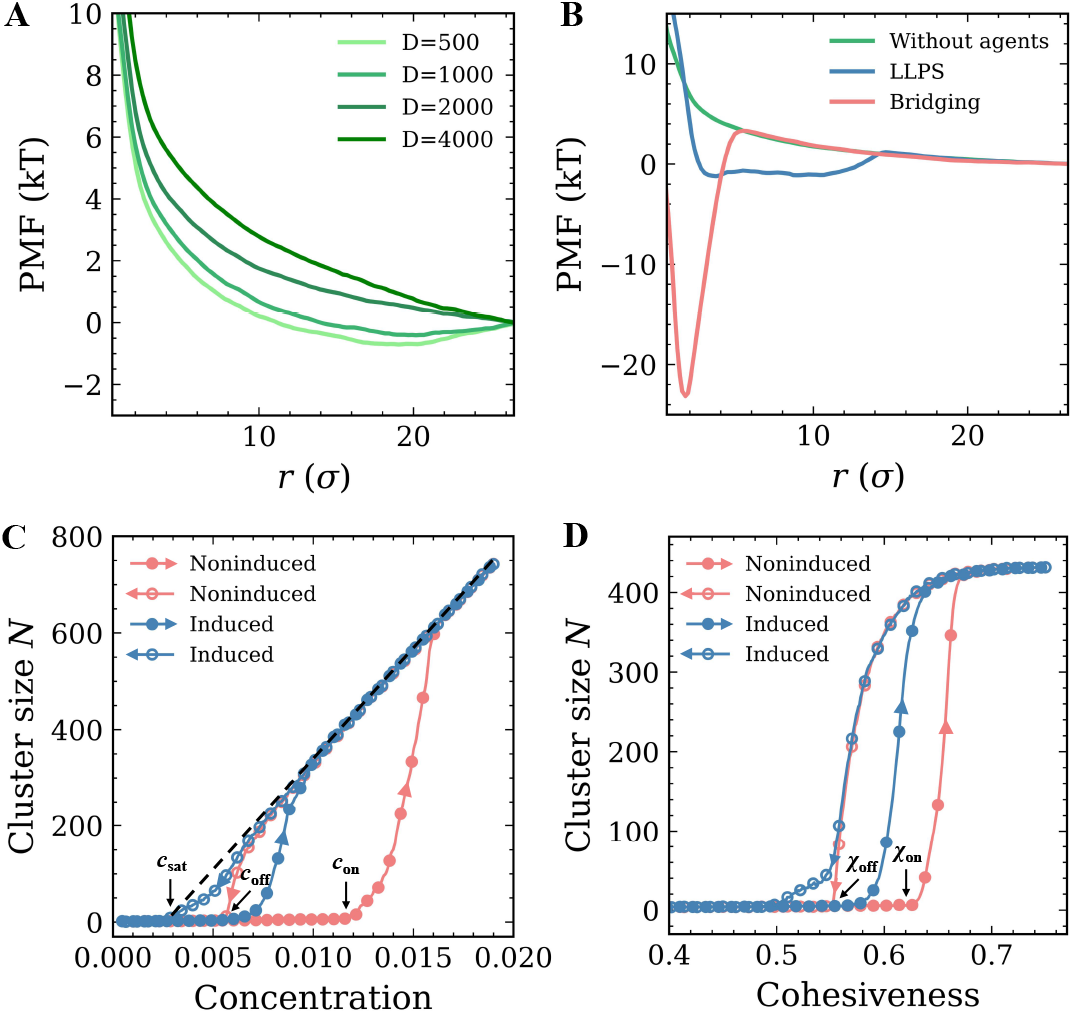
PMF and hysteresis calculations show key thermodynamic properties of chromatin, CREs and the agents. (A) PMF profiles of two-body CRE interaction with varying contour distances (D=500, 1000, 2000 and 4000 beads), in the absence of agents. For simplicity, the same initial spatial distance is used in all the cases, which does not correspond to the minima of the PMFs. (B) Comparison between the effects of agent LLPS and bridging on the PMF profiles of two-body CRE interaction. The case of no agents is shown as a reference. The initial contour distance between the CREs is D = 2000 and the agent concentration is *c* = 0.01 in all the cases. (C) The number of agents in the largest cluster as a function of the agent concentration in the LLPS hysteresis simulations with (blue) and without (red) the nucleation stimuli from the CREs. A constant agent cohesiveness *χ* = 0.6 is used. The symbols and the arrows of the lines both show the direction of the concentration change. The corresponding positions of *c*_sat_, *c*_off_ and *c*_on_ are marked. (D) Hysteresis profiles of LLPS at varying agent cohesiveness and constant agent concentration *c* = 0.01. The corresponding positions of *χ*_off_ and *χ*_on_ are marked. All the hysteresis data are averaged over 100 independent simulations.

Before interrogating the different effects of LLPS and bridging on multi-way genomic interaction, it is instructive to first comprehend the phase diagram of agent condensate in our system. We first study the phase behavior of the system as a function of the total agent concentration *c* with the agent cohesiveness strength *χ* being fixed. To efficiently do that, we carry out hysteresis simulation with two opposing periods. In the first half of the simulation, we increase the agent concentration in a slow and linear fashion from a dilute baseline to a level that is dense enough to generate a large droplet. We then reverse the process so that the droplet shrinks and dissolves at a critical size. During the simulation, we analyze the size of the largest agent cluster as a function of the concentration. Since the onset and dissolution of the droplet is stochastic, we average over 100 independent simulations to get the ensemble-averaged hysteresis curve as plotted in Figure 2C. The simulations clearly show that the droplet has an onset concentration *c*_on_ that is higher than its dissolution concentration *c*_off_, a sign of hysteresis. For a stable droplet, its linear size dependence on the agent concentration is a consequence of the dilute phase concentration being constant, a hallmark of homogeneous LLPS. This constant is often referred to as saturation concentration *c*_sat_ or critical concentration. *c*_sat_ is smaller than *c*_off_ because the latter is the total concentration of the agents including the contribution from the critical droplet that is not negligible in our closed finite system. Of note, chromatin with inactive CREs is included in our hysteresis simulation and it can enhance LLPS through crowding. We then turn on the CREs to study how nucleation shapes the hysteresis of LLPS. As shown in Figure 2C, CREs can induce the LLPS at a concentration in between *c*_off_ and *c*_on_. On the other hand, the nucleation sites can stabilize the agent condensate against dilution. Below *c*_off_, bridging takes over LLPS as the driving force to cluster up the CREs. Similarly, in the 1D phase diagram as a function of the agent cohesiveness strength *χ* for a given agent concentration (Figure 2D), one can see that induced LLPS occurs in the regime between the dissolution cohesiveness strength *χ*_off_ and the onset cohesiveness strength *χ*_on_, and bridging happens below *χ*_off_. The individual hysteresis curves of the agent condensates, the distribution of the critical droplet size, and the averaged hysteresis curves of the CRE hubs are shown in Figure S3-8 as reference.

In the above hysteresis simulations, the variables are slowly changing so that the system evolves in a quasi-static manner, which is suitable for calculating the thermodynamic phase diagrams. We now carry out nonequilibrium simulations to investigate the dynamics of the higher-order catalyzing system, with the focus on how the agents stabilize the transition state of the higher-order reaction, i.e., the CRE hub. We simulate the nonequilibrium dynamics of the activated CREs in systems with varying total concentrations of the agents. Figure 3A shows the final-state snapshots from simulations at three typical agent concentrations, which correspond to the scenarios of single phase (no droplet), small droplet, and large droplet. In the first scenario, the 5 CREs are well separated in space. In the latter two cases, with a transparent visualization of the droplets, one can clearly see that all the 5 CREs are brought together by the agent condensation. We analyze the time-dependent clustering of the CREs at varying agent concentrations and average the results over a set of independent simulations (typical individual clustering profiles can be found in Figure S9). The ensemble-averaged profiles are plotted in Figure 3B, which reveals a remarkable acceleration effect of the agents on the clustering of the CREs. In a reference system, we set the CREs free by taking them off the chromatin polymer. As shown in the dash line of Figure 3B, these free “CREs” can almost instantly form a hub upon agent condensation, in comparison to the chromatin-constrained cases. This result demonstrates that, adversary to the catalytic effect of the agent condensation, chromatin slows down the clustering of the CREs. Our simulations predict that the averaged cluster number of the CREs decays exponentially to 1 as all the CREs cluster up (Figure S10). This is in line with our higher-order catalysis model and extracting the exponential decay constant gives us the activation rate constant in Equation 2. The activation rate constant as a function of agent concentration is shown in Figure 3C, which looks like a ReLU (Rectified Linear Unit) activation function^43^. Namely, above an activation threshold, further increasing the agent concentration accelerates the CRE clustering in an approximate linear way. The threshold is around the onset concentration of induced LLPS, indicating that LLPS is the underlying driving force of the CRE clustering (maturation of the condensates and the rate constant as a function of the matured condensate size can be found in Figure S11, S12). The positive correlation between the agent concentration and the accelerating effect of the LLPS is analogous to the concentration-dependent enzyme activity. This interesting similarity, again, supports our hypothesis that LLPS can serve as a higher-order catalyst or super-enzyme in gene regulation. However, it is worth noting that, while a higher concentration of enzymes speeds up the reaction rate by acting on more substrates, increasing the agent concentration allows the LLPS to escalate the intrinsic rate constant of the higher-order reaction. An upper limit of the agent concentration is the spinodal point, above which LLPS will spontaneously happen. One should be reminded that the activation rate constant describes the average speed of the higher-order reaction over a population of cells. To appreciate the stochastic nature of the activation process, we analyze the first-passage time (FPT) of the CRE clustering, i.e., the time when the hub of all the CREs first appears. The results of three different conditions are plotted in Figure 3D, which shows that the distribution of the FPTs gets wider when the clustering slows down. In the above simulations, we have used a relatively weak attraction strength between the CREs and the agents. Under such a condition, we observe that successful nucleation of LLPS is often facilitated by the synergy of two CREs that diffuse close to each other in space. The induced droplet then grows and recruits the other CREs to form the CRE hub (Figure S13). In the case of stronger nucleation strength, induced LLPS is observed to happen simultaneously at multiple CREs and the formation of the CRE hub is accompanied by droplet coalescence and Ostwald ripening (Figure S14).

**Figure 3.**
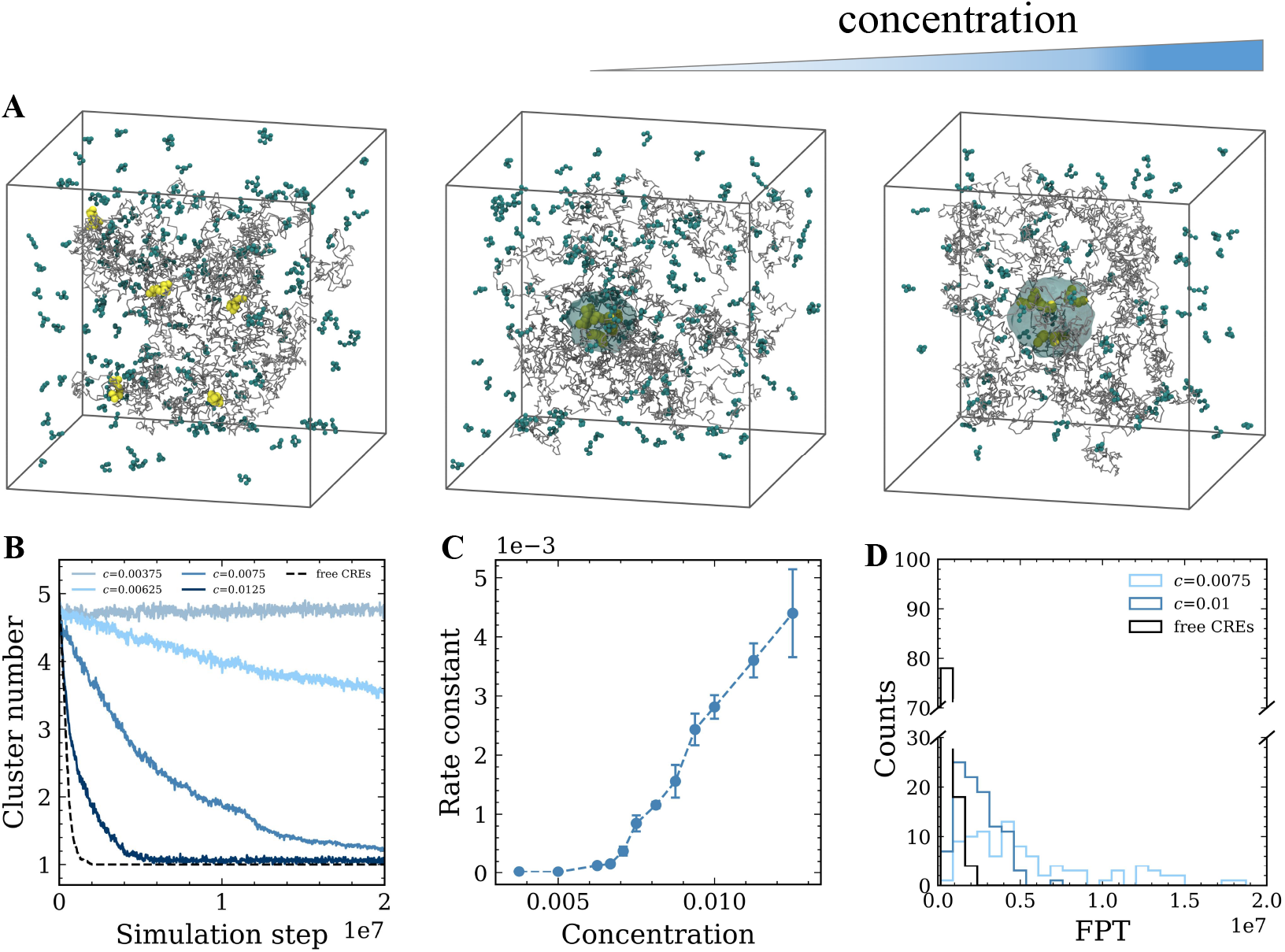
Nonequilibrium simulations and analyses of the CRE clustering driven by cohesive agents. (A) Typical simulation configurations at ascending total concentration of the agent. For visual clarity, the largest agent cluster associated with the CRE hub is shown with a transparent droplet-like style. The chromatin chain is shown in gray, CREs in yellow, and agents in cyan. (B) Time-dependent CRE clustering profiles at different agent concentrations (averaged over 100 independent simulations). The clustering profile of free CREs (*c* = 0.01) is presented in the dashed line as a reference. (C) Rate constant of the CRE clustering as a function of the agent concentration based on the exponential regression of the averaged clustering profiles. (D) FPT analysis shows great stochasticity of the CRE clustering. A constant agent cohesiveness *χ* = 0.6 is used in all the nonequilibrium simulations.

The above nonequilibrium simulations demonstrate the ability of LLPS to accelerate the clustering of the CREs. On the other hand, we have not observed bridging-based activation of the CRE hub. One possible reason is that the employed CRE-agent attraction that is strong enough to nucleate LLPS could be too weak to elicit bridging. To gain deeper understanding of this non-LLPS mechanism, we extend our nonequilibrium simulation to a strong bridging system in which the effective two-body avidity between the CREs can be higher than 20kT due to the physical crosslinking by the agents (Figure 2B). We tune down the agent-agent cohesiveness to make sure that LLPS is inhibited. Figure 4A shows the bridging-driven clustering profile of the 5 CREs as a function of time, compared against a LLPS case with the same agent concentration. These ensemble-averaged simulation results reveal fundamental differences of the clustering dynamics between the two distinct molecular mechanisms. Unlike the case of LLPS, after we turn on the bridging interaction, the time-dependent cluster number of the CREs does not follow an exponential decay. It is interesting that the decay driven by bridging is fast at the beginning, even surpassing the decay speed under LLPS, but slows down as it gets close to 1. Such clustering dynamics indicates that strong bridging can rapidly pair up two CREs but is not as effective in mediating multi-way genomic contacts.

**Figure 4.**
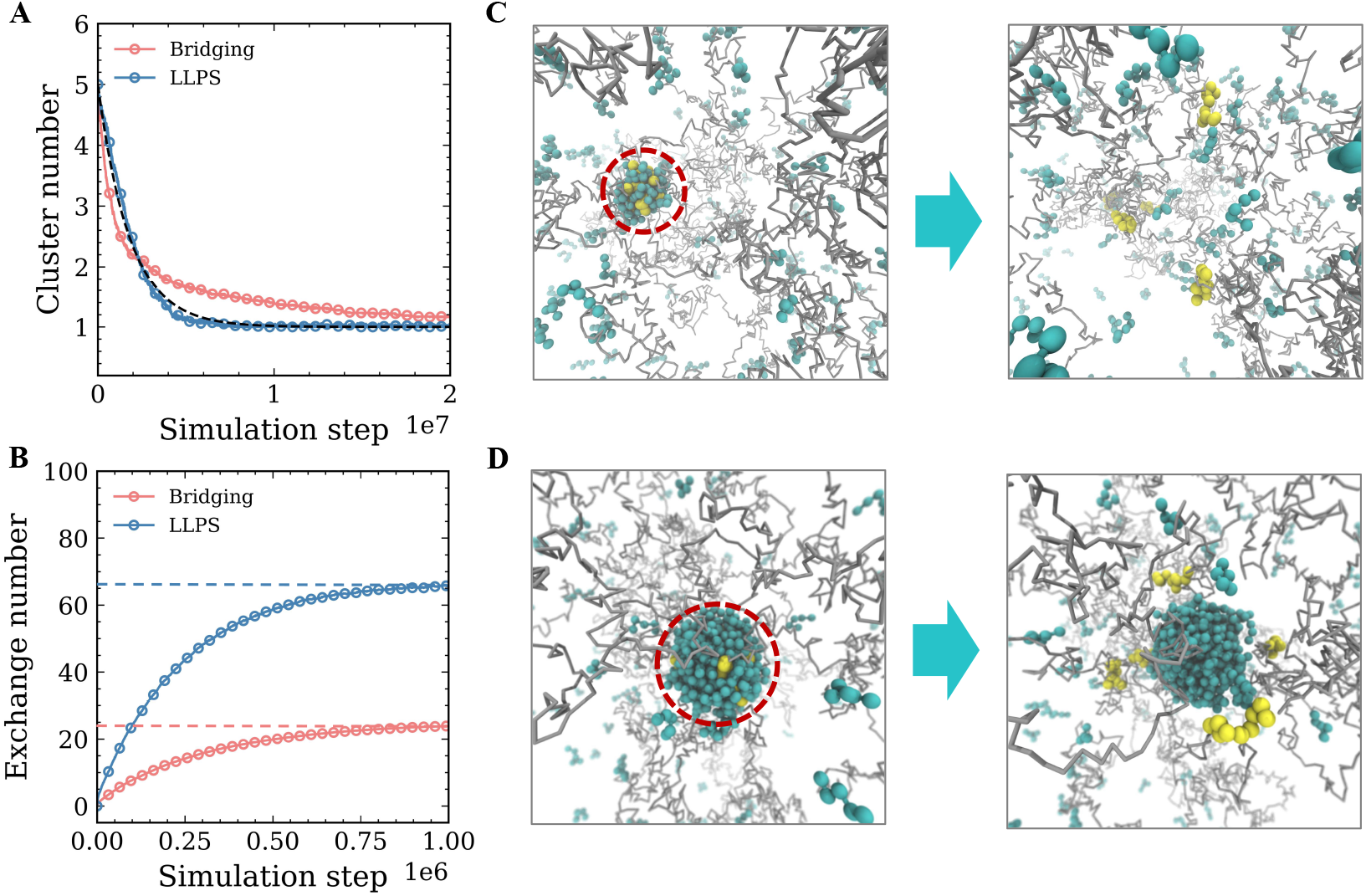
Comparison between bridging and LLPS as different driving forces of CRE clustering. (A) Time-dependent CRE clustering profiles under agent bridging and LLPS (averaged over 100 independent simulations). Exponential regression only applies to the LLPS case as shown in the dashed line. (B) Averaged exchange numbers of agents as a function of time in the equilibrated CRE-agent condensates for the bridging and LLPS systems, the dotted lines indicate the asymptotic values predicted based on the sizes of the condensates. (C) Simulation configurations of the bridging system before and after the CRE-agent interactions are turned off. The condensate marked in the red circle disappears. (D) Simulation configurations of the LLPS system before and after the CRE-agent interactions are turned off. The condensate marked in the red circle persists.

Another important difference between LLPS and bridging we discovered is that the latter is not sensitive to the agent concentration (Figure S15). Across the wide concentration range we tested, bridging fails to be a robust mechanism to drive the formation of the CRE hub. Of note, once formed, the bridging-based condensate of the CREs and agents is thermodynamically stable. However, it is different from the LLPS-based condensate in many ways. First, it is tiny in size, even smaller than the critical droplet formed via LLPS. Second, its mass exchange with the environment is slower than that of the LLPS-based droplet. To see that, we analyze the time correlation of the agent constituents for both bridging-based and LLPS-based condensates. Despite the larger back-flux of agents into the LLPS-based condensate in our confined system, the net depletion of agents is still much faster with LLPS, in comparison to bridging (Figure 4B). Finally, the CRE-agent affinity is required to stabilize the bridging-based condensate but not the droplet formed via LLPS. In other words, dismissing the CRE-agent affinity will dissolve the bridging-based condensate (Figure 4C), whereas the LLPS-based droplet, once induced, is self-sustainable without the nucleation sites (Figure 4D). Additional work or energy input is needed to dissolve the liquid droplet in the phase-separated system. It is a pitfall to think that turning off the nucleation sites alone can reverse the LLPS. As proposed in our higher-order catalysis model, to serve as a super-enzyme, LLPS not only needs to selectively respond to nucleation stimuli but also should be able to recognize specific downstream reactions as resetting cues. Our systematic comparison between bridging and LLPS highlights the advantage of LLPS in mediating long-range multi-way interactions of chromatin and stresses its thermodynamic irreversibility that calls for specific dissolution mechanisms.

## Discussion and conclusion

A growing body of research suggests that LLPS is involved in regulating many nuclear activities^44^, with the most intensively studied being transcription^45,46^. However, it remains elusive whether LLPS is required for Pol II transcription^16,17^. Some key features of the CRE hub and its associating condensate seem to be at odds with the canonical understanding of LLPS-driven MLOs. To resolve this paradox, we propose the higher-order catalysis model that integrates chromatin mechanics, LLPS and chemical reactions in a unified theoretical framework. Here, we discuss the implications of this model.

It is well known that transcription occurs in bursts or pulses^47^, and transcription can be regulated by changing the bursting frequency or amplitude. In the framework of higher-order catalysis, the bursting frequency as the reaction rate can be modulated by the LLPS as the super-enzyme, whereas the bursting amplitude as the amount of product can be controlled by the number and strength of the CREs as the reactants. Of note, being a super-enzyme means that LLPS can significantly expedite the higher-order reaction, but it is not indispensable for transcription^17^. Nevertheless, we expect that for some cell-specific genes, the chromatin reorganization energy could be so large that higher-order catalysis is required to activate the transcription. While the higher-order catalysis model provides a mechanistic explanation for the regulation of transcriptional bursting, its application would depend on the type of the gene. For genes that do not or no longer require multiple enhancers, such as housekeeping genes and some highly active cell-specific genes, their expression is likely regulated by other mechanisms. How different modes of transcription regulation interplay to orchestrate layered control of gene expression is an important question for future investigation.

Our model highlights an underappreciated function of the seemingly unnecessarily long genomic distances between CREs and their targets, that is to constitute an activation barrier to inhibit aberrant gene expression. In this light, a stretch of purely inert genome that is noncoding, inactive, and void of protein binding sites should not be considered as junk DNA^48^ as it can shape the free energy landscape of the higher-order genomic reaction via changing the conformational entropy of chromatin. If there is no bottleneck in the higher-order reaction to limit its reaction rate in the first place, a catalytic machinery would be meaningless. Our simulation suggests that chromatin looping that mediates only two-body contact is not a very efficient way to construct a strong activation barrier (Figure 2A). Compared to two-body contact, multi-way contact is a good strategy for chromatin to significantly holster the barrier level, as the probability for multiple CREs to simultaneously encounter each other in space against long genomic distances is orders of magnitudes lower. On the other hand, decision making based on the information encoded in multiple CREs also allows for more robust and cell-specific regulation of transcription and alternative splicing. Taking these advantages together, it is becoming more comprehensible why mammalian cells bother to have long-range multi-way genomic interactions.

Our computational comparison between LLPS and bridging suggests that long-range weak attraction is more advantageous than short-range strong attraction in driving multi-way long-range genomic contacts. This is because the repulsion between the CREs is due to the long-range entropic force of chromatin, and such a soft barrier can be better counteracted by long-range weak attraction. Another advantage of LLPS over bridging is its sensitivity to the agent concentration, which allows dynamic regulation and feedback control of the catalytic activity of LLPS, since some agents can be the product of the higher-order reaction they regulate. For simplicity, our minimal computational model considers only one type of agent and is therefore limited to homogeneous LLPS. In reality, the nucleus is a pool of different agents that can undergo a wide spectrum of heterogeneous LLPS in response to varying nucleation stimuli. This can lead to a family of LLPS-based super-enzymes that are ready to catalyze different higher-order reactions. While induced heterogeneous LLPS offers good catalytic specificity and diversity, sharing a subset of core agents could further allow intricate interplay between different super-enzymes and simultaneous catalytic regulation at the global scale.

It is well known that the rate of an enzymatic reaction depends not only on the enzyme concentration, but also on temperature^49^. Modest increase of temperature that does not denature the enzymes in general speeds up enzymatic reactions. However, such temperature dependence does not necessarily apply to the LLPS-based higher-order catalysis. One important thermodynamic implication of our model is that the chromatin-based reaction barrier and the LLPS-based super-enzyme have opposite responses to temperature change. Polymer physics predicts that the entropic reaction barrier due to chromatin reorganization will strengthen as temperature increases. In contrast, raising temperature would weaken the LLPS of agents as it is enthalpy-driven at the cost of translational entropy. In combination, these two factors tend to synergistically lower the rate of higher-order reaction at higher temperature. Parallel to active temperature-responsive regulation programs, such passive thermodynamic effects could have implications in early embryonic development and immune response^50,51^.

The higher-order catalysis is associated with energy transformation. At the beginning of the enzymatic cycle, thermodynamic free energy is stored in the metastable single-phase state of the agents. Induced by the CREs, the phase transition then uses the free energy to reorganize the chromatin segment into multi-loops. The reorganized chromatin that mediates the multi-way CRE contact has a lower entropy, and higher free energy. To complete the enzymatic cycle, the agents need to be reset to the single-phase state. Since LLPS is a spontaneous process that does mechanical work, the dissolution of the droplet requires energy input or external work so that the higher-order catalyst is not a perpetual motion machine. Such energy can be provided by the downstream chemical reactions triggered by the CRE clustering. While the LLPS does mechanical work to stabilize the transition state during the enzymatic cycle, it is the higher-order reaction itself that pays the free energy. This makes the cycled phase transitions distinct from biomolecular motors that require external energy sources. Unlike the Cohesin-driven loop extrusion^2,3^, the LLPS-based catalytic machinery does not consume chemical energy stored in ATPs to do mechanical work. It is worth noting that the forms of mechanical work done in the two processes are also different. During the continuous extrusion of chromatin, Cohesin can sense the genomic details along the chromatin contour. In contrast, during the LLPS-driven condensation, the agents selectively act on the CREs so that the genomic contents between the CREs have limited effect on the CRE clustering. The diversity of working mechanisms and energy sources would be beneficial for robust gene regulation.

We did not explicitly model the dissolution process because it can be triggered by different molecular mechanisms depending on the nature of the higher-order reaction. Although we have so far focused on transcription regulation as an example, the model of higher-order catalysis is transferrable to other genomic events such as gene silencing and DNA replication. Super-resolution microscopy has revealed that Polycomb-group proteins for gene repression form many nanoscale condensates^52^. It has been recently reported that these silencing Polycomb condensates promote epigenetic marks but are dispensable for sustained chromatin compaction^53^. Although more details are needed, the Polycomb condensation is likely driven by LLPS induced by repressive CREs whose multi-way contact activates gene silencing. Once the rate-limiting transition state is stabilized by the LLPS as higher-order catalyst, the subsequent spreading of epigenetic marks and compaction state can proceed with a relatively high speed. In the case of DNA replication, the transition state can be the hub of multiple replication origins^54^. By driving the formation of such replication origin hubs, LLPS can promote the downstream spreading of replication. In general, the higher-order catalysis model suggests that chromatin reorganization is a universal activation barrier and cycled phase transition a universal enzymatic machinery for varying genomic programs. We emphasize that the role of LLPS in our model is to accelerate rather than compartmentalize the higher-order reactions. After the reaction activation, whether the producing stage of transcription, epigenetic marking or DNA replication occur in specific factories are important questions but out of the scope of this paper.

We expect the higher-order catalysis mechanism to have profound impacts on the higher-order structure and dynamics of chromatin. With the textbook picture of chromatin folding being outdated, it is becoming more accepted that the genome is organized into a hierarchy of compartments despite the debate on whether such compartmentalization starts at the nucleosome level or the controversial 30 nm fiber level^35,55^. The 3D structure of the compartments at the single-cell level is under intensive study both experimentally and theoretically^56,57^. Our recent structural model predicts that chromatin folds into a series of treelike topological domains connected by an open backbone in single cells^32,33^. The tree domains house abundant multi-way contacts which is puzzling from a polymer physics point of view. It is natural to ask how these many multi-way contacts could possibly arise and what are their potential functions. The higher-order catalysis model fills the gap between higher-order chromatin structure and genomic function, and further predicts that most multi-way contacts are highly transient. They are not passive byproducts of chromatin activities but actively formed and timely deconstructed for specific genomic functions. According to equilibrium thermodynamics and polymer physics, rich multi-way interactions will strengthen the crosslinking of chromatin and turn it into a hydrogel that is a state of solid. However, in the picture of higher-order catalysis, the multi-way contacts come and go quickly, rearranging chromatin constantly. Such nonequilibrium activities allow chromatin to stay in a highly dynamic, liquid-like state, which is consistent with recent experimental observations^58^.

While the LLPS-based super enzymes can heavily shape the dynamic structure of chromatin, the genomic architecture as the context of the higher-order reaction can in return significantly influence the higher-order catalysis. Although we have only simulated homogeneous space for simplicity, our theoretical model can be applied to heterogeneous nuclear environments. As a heteropolymer, chromatin undergoes polymer-polymer phase separation (PPPS) at the megabase scale that leads to the A/B compartmentalization^34,50,56,59^. The two types of compartments differ in physicochemical properties such as DNA packing density, solvent quality, and molecular diffusion constant. These factors will all affect the LLPS and the clustering of CREs. Moreover, there are many membrane-less nuclear bodies of varying functions inside the nucleus which create a considerable amount of interface with chromatin^44^. Akin to heterogeneous catalysis, interfaces between chromatin compartments and the nuclear bodies could facilitate the clustering of specific CREs driven by interfacial LLPS, as shown in Figure 5. Wetting properties on these 2D surfaces and even on the 1D chromatin fiber might play important roles in such heterogeneous higher-order catalysis.

**Figure 5.**
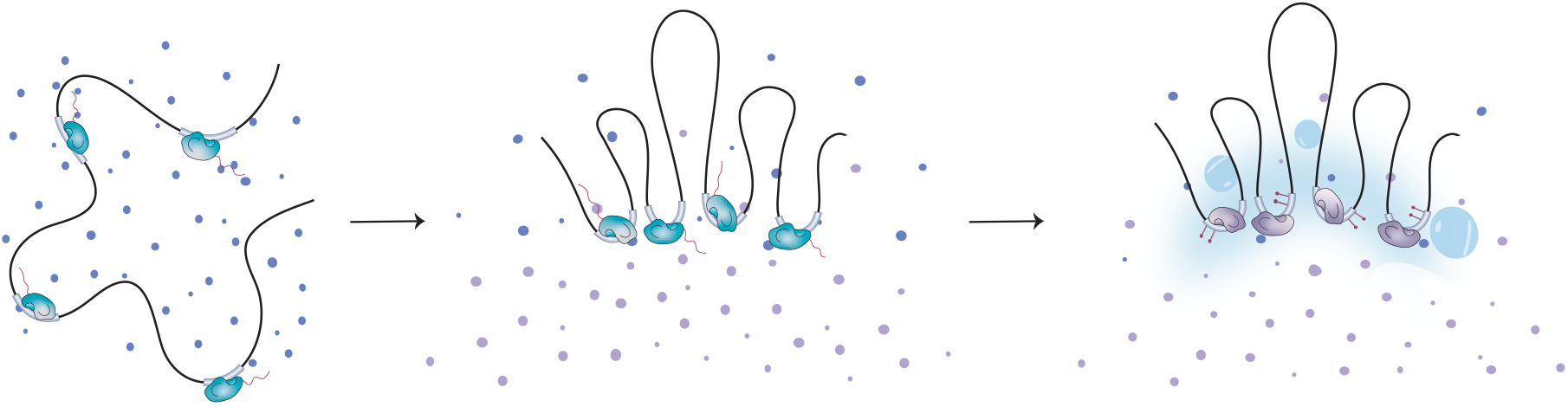
Schematic illustration of the heterogeneous higher-order catalysis. Same as the flow of Figure 1, but here the higher-order reaction occurs on a liquid-liquid interface.

One fascinating characteristic of chromatin is fractal, which is a mathematical concept that means self-similarity across different scales^32^. The fractal structure of the genome is manifested in its hierarchical compartmentalization^36^. The high-order catalysis model suggests that the concept of self-similarity can be extended from chromatin structure to its biochemical function. It shows that chromatin-mediated biochemical reactions and their catalysis are hierarchically organized at different scales. In particular, the activation of CREs is realized by biochemical reactions catalyzed by specific enzymes. The activated CREs then catalyze the nucleation of the LLPS of agents. The induced LLPS further catalyzes the higher-order reaction at a larger scale. Of note, while catalyzed biological events happen at multiple scales, their catalyzing mechanisms are not the same. Self-similarity is often associated with criticality of the system. Our model predicts that the agent concentration is maintained in between the critical point and the spinodal point to take advantage of the metastability of the single phase. This regime allows the LLPS to be sensitive to the cues of nucleation and dissolution. If phase transition is so important in gene regulation as predicted, the nucleation process of itself needs to be highly regulated. Using ncRNAs and their transcription is a good strategy to achieve such hierarchical regulation. Therefore, it is not surprising that Pol II has an intrinsically disordered region that facilitates LLPS^60^, and that noncoding transcription is implicated in gene silencing^59^. Mediating multi-way genomic interaction through the condensation of agents rescales the cutoff length of contact to the size of the regulatory condensate. From a fractal point of view, the induced LLPS increases the length unit of the measurement, which is equivalent to shortening the contour length of chromatin. In a sense, during the higher-order catalysis, chromatin renormalizes itself to reduce the activation barrier. These insights suggest that self-similarity is a pivotal theme in chromatin biology. It is worth noting that, given the hierarchical framework, LLPS can play duo rules without conflict at different scales in gene regulation. The regulatory condensate can be viewed as MLO at a small spatiotemporal scale to hold molecular reactions, but a transition state at the larger scale of higher-order catalysis.

In conclusion, the higher-order catalysis model reveals a new relation between the physical chemistry of LLPS and its biological function. The model provides explanations to the abundance of multi-way genomic contacts, the long-range communication of multiple CREs, and the life cycle of CRE-associated condensates, that are puzzling in the conventional paradigm of polymer physics and LLPS-based macroscopic MLOs. In biological systems, enzymes are ubiquitous and needed in almost all the biochemical reactions. Our work generalizes the concept of enzyme from single molecules to the phase behaviors of nonstoichiometric supramolecular assemblies. The CRE-induced nucleation of specific LLPS, the agent condensation that aggregates the CREs, and the dissolution of the droplet constitute a higher-order enzymatic cycle that acts on the higher-order structure of chromatin as its substrate. The new role of LLPS as a super-enzyme can be implicated in a diversity of chromatin-mediated higher-order reactions such as gene expression, silencing, epigenetic marking, and DNA replication. Our work leads to a hierarchical picture of catalyzed genomic reactions and calls for efforts to target catalytic LLPS in therapeutic development especially for neurodegenerative diseases and cancer. The higher-order catalysis model provides mechanistic support for our recently developed 3D forest model of chromatin^32^. Together, these two models self-consistently provide holistic insights on the structure and dynamics of chromatin as a complex living polymer, and how it exploits LLPS, a thermodynamic driving force from nature, to regulate itself. A strange yet marvelous view of chromatin folding emerges, in which the genome is organized into a four-dimensional (4D) “forest” of tree-like domains that are topologically variable and dynamic in single cells. Cycled LLPS leads to transient condensates like the lights of firebugs in the woods, except for that their firing and fading constantly reorganize the 4D chromatin forest. We hope the higher-order catalysis framework provides a new lens through which intriguing multiscale biological phenomena can be better understood.

## Acknowledgements

We acknowledge financial support from the Shenzhen Bay Laboratory Open Fund Project (SZBL2021080601013), and support from the Shenzhen Bay Laboratory Supercomputing Center. We thank Xiangli Liu for the schematic illustration of the model.

