## Supporting Information for "Phase separation as higher-order catalyst"

### 1. Simulation methods

In this work, a bead-spring polymer model is introduced to investigate the interactions between the CREs of chromatin and the trans-acting agents. The chromatin fiber is modeled as a polymer chain with 5000 coarse-grained beads, 5 CREs (each consisted of 10 consecutive beads) separated by 1000 polymer beads (defined as distance D = 1000) are distributed on it. Agents are represented by 5 consecutive beads, which can provide multivalent interactions to crosslink distant CREs. All beads have mass $m = 1$ and radius $\sigma = 1$. Bonds between adjacent beads are described by a harmonic potential that is, $U_b(r) = k(r - r_0)^2$, where $k$ denotes the spring constant and is set to 50, $r_0$ is the equilibrium bond distance and is set to 1 $\sigma$. The non-bonded isotropic interactions between all types of beads are modeled using the standard 12-6 Lennard-Jones (LJ) potential that is, $U_{LJ}(r) = 4\epsilon\left[\left(\frac{\sigma}{r}\right)^{12} - \left(\frac{\sigma}{r}\right)^{6}\right]$, for all $r < r_c$, where $r_c$ refers to the cutoff distance beyond which the non-bonded interactions are neglected. The LJ potential is truncated at $r_c = 2.5\ \sigma$. In our simulations, the effects of induced phase separation and bridging on CRE clustering are considered, the interaction parameters are listed in Table 1.

**Table 1**. LJ interaction parameters.

| | LLPS system | Bridging system |
|---|---|---|
| Chromatin - chromatin | 0.2ϵ | 0.2ϵ |
| CRE - CRE | 0.2ϵ | 0.2ϵ |
| Agent – agent | 0.6ϵ | 0.2ϵ |
| Chromatin – CRE | 0.2ϵ | 0.2ϵ |
| Chromatin - agent | 0.2ϵ | 0.2ϵ |
| CRE - agent | 0.74/0.85ϵ | 1.4ϵ |

Steered molecular dynamics (SMD) simulations are conducted to estimate the free energy profile of CRE clustering in two cases: (i) the clustering of two CREs with varying contour distances (D = 500, 1000, 2000 and 4000) on the chromatin chain without agents; (ii) the clustering of two CREs on the chromatin chain driven by induced phase separation and bridging respectively, the contour distance between the two CREs is fixed at D = 2000 and the agent concentration $c$ is set to 0.01. In SMD simulations, a guiding potential is exerted on the two CREs to bring them together at a constant velocity. The potential of mean force (PMF) profile is then estimated along the reaction coordinate by calculating work and employing Jarzynski's equality(*1*). Here, the reaction coordinate is defined as the physical distance between the two CREs.

To understand the phase behaviors of the agents, concentration-scanning and cohesiveness-scanning hysteresis simulations are conducted, respectively. In both cases, the chromatin chain without or with CREs (denoting non-induced and induced phase separation) are considered. In the concentration hysteresis simulation, the agent concentration $c$ first gradually increases from 0.0004 to 0.019 through agent addition, and then it is gradually decreased to 0.0004 through agent deletion. The $\epsilon_{\text{agent-agent}}$ and $\epsilon_{\text{CRE-agent}}$ are fixed at 0.6 kT and 0.74 kT, respectively, other interaction parameters are set to 0.2 kT. At each add/delete interval, $2\times10^5$ simulation steps are performed; In the cohesiveness hysteresis simulation, the $\epsilon_{\text{agent-agent}}$ first gradually increases from 0.35 kT to 0.75 kT within $2\times10^7$ simulation steps, and then it is decreased gradually to 0.35 kT within another $2\times10^7$ simulation steps. The $\epsilon_{\text{CRE-agent}}$ is fixed at 0.74 kT, other interaction parameters are set to 0.2 kT, the agent concentration $c$ is kept at 0.01. In both cases, the largest cluster size of agents is monitored to assess the hysteretic phase separation behaviors.

Regarding to the induced phase separation, we further investigate the effect of agent concentration on CRE clustering dynamics in more details, in which $c$ is ranged from 0.00375 to 0.0125. The clustering of free CREs without polymer restriction at $c = 0.01$ is studied as a reference. Besides, another simulation, in which the $\epsilon_{\text{CRE-agent}}$ is tuned to 0.85 kT, is performed to investigate the effect of interaction strength between CREs and agents on CRE clustering dynamics. The $\epsilon_{\text{CRE-agent}}$ and $\epsilon_{\text{agent-agent}}$ are changed to 1.4 kT and 0.2 kT, respectively, to study the CRE clustering behaviors driven by the bridging mechanism with different agent concentrations ($c$ = 0.00375, 0.00625, 0.075, 0.01). For all simulation systems, the initial configuration first undergo $2\times10^5$ steps of energy minimization with all interactions turned off. After then, the interactions are turned on and $2\times10^7$ simulation steps are performed to make the system equilibrated. Trajectories are saved each 5000 steps for outcome analysis.

The initial configuration of the chromatin chain is generated by self-avoiding random walk (SAW), agents are uniformly distributed in the box as the simulation starts. The temperature of the simulation system is controlled by Langevin thermostat. The simulation parameters are set as follows: damping parameter $\gamma$ = 100, temperature T = 1, and integration time step dt = 0.01, expressed in dimensionless units. All simulations are conducted using the Large-scale Atomic/Molecular Massively Parallel Simulator (LAMMPS) package(*2*). The Visual Molecular Dynamics (VMD) software(*3*) is used for structure visualization and snapshot production. Analyses of simulation trajectories are carried out with in-house scripts. To ensure the adequacy of conformational sampling, 100 independent runs are performed and averaged for each simulation system.

### 2. PMF analysis

As shown in Figure S1, there is a free energy barrier (about 1 kT) for the induced phase separation system at r = 15σ, where a small agent cluster formed on one of the two approaching CREs. This small energy barrier can be overcome via thermal fluctuation, which indicates that the CRE clustering can proceed spontaneously in this case. After crossing the energy barrier, the PMF profile decreases and exhibits a wide and shallow free energy basin. Such long-range attraction mediated by the LLPS of the agents can facilitate the formation of CRE hub when it comes to many-body CRE interactions. For the bridging system, a free energy barrier (about 2 kT) appears at $r = 5\sigma$, which denotes that the interaction range is much shorter than that of the induced phase separation system. The PMF profile sharply decreases and shows a narrow valley at distance $r$ = 1.8σ after the energy barrier has been crossed, the final configuration shows an establishment of agent bridges between the two CREs, as depicted in Figure S2.

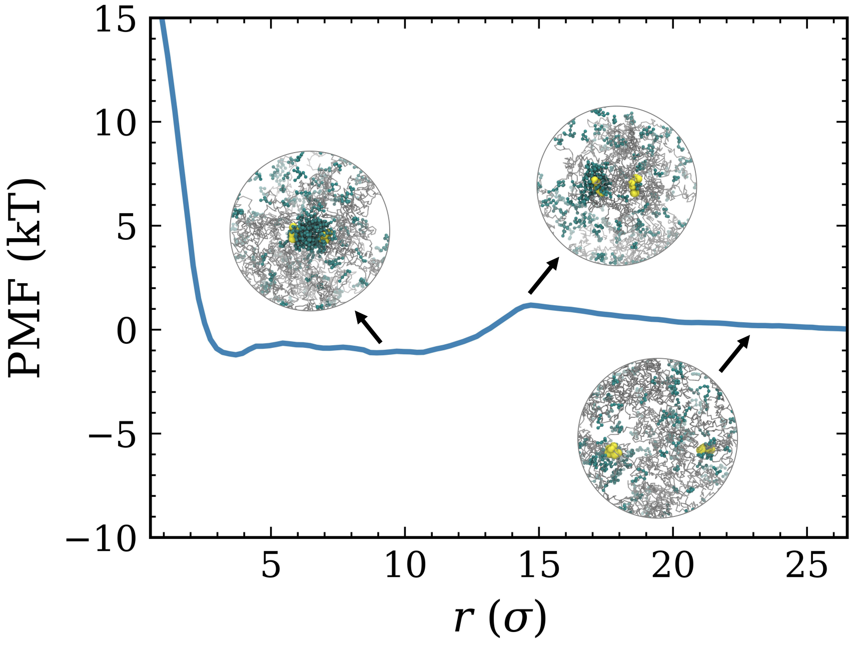


**Figure S1.** PMF profile of CRE interaction as a function of the distance between two CREs for the induced LLPS system at agent concentration $c = 0.01$. Insets are typical snapshots during the pulling process.

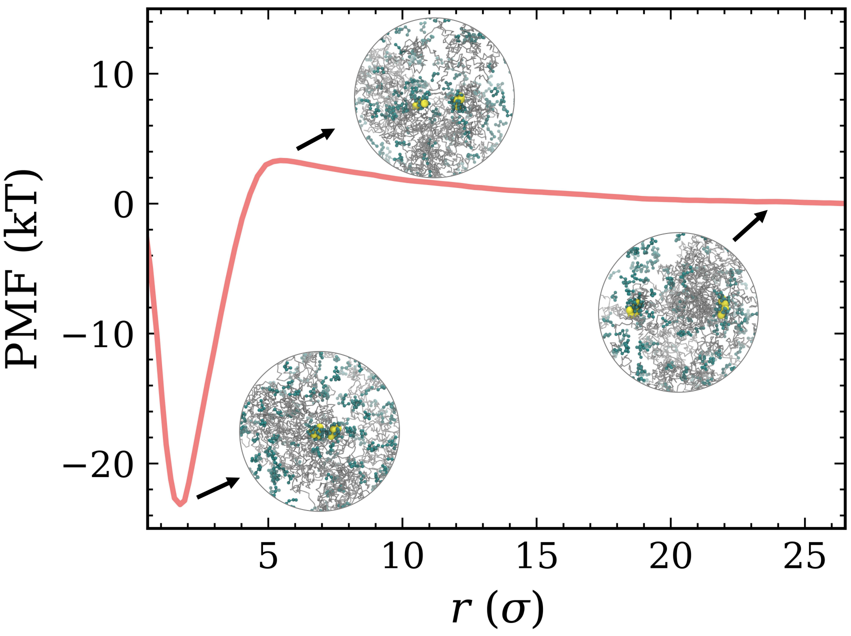


**Figure S2.** PMF profile of CRE interaction as a function of the distance between two CREs for the bridging system at agent concentration $c = 0.01$. Insets are typical snapshots during the pulling process.

### 3. Hysteresis profiles of agent phase transitions

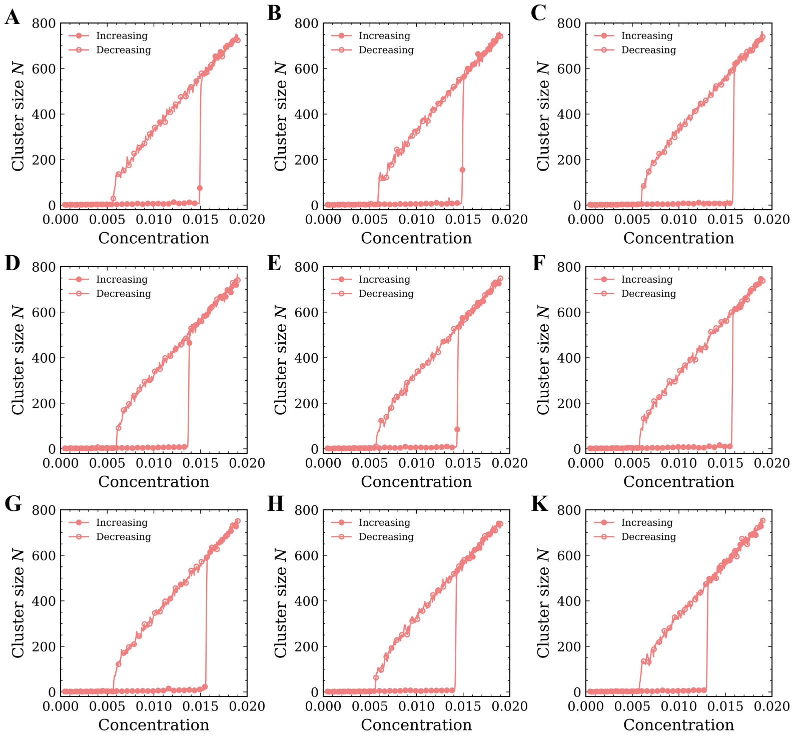


**Figure S3.** The cluster size of agents as a function of agent concentration with the induction of CREs turned off in several independent runs.

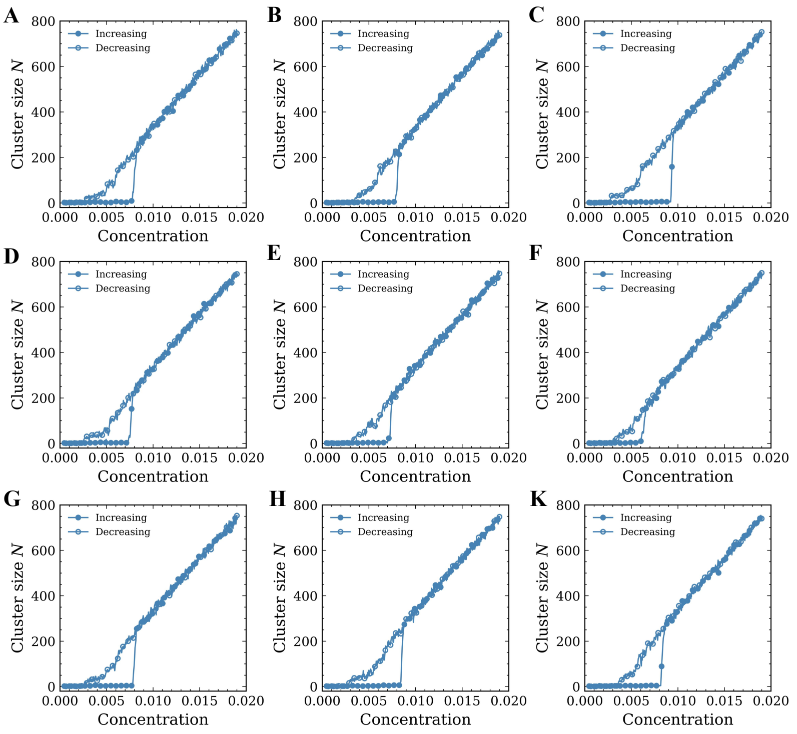


**Figure S4.** The cluster size of agents as a function of agent concentration with the induction of CREs turned on in several independent runs.

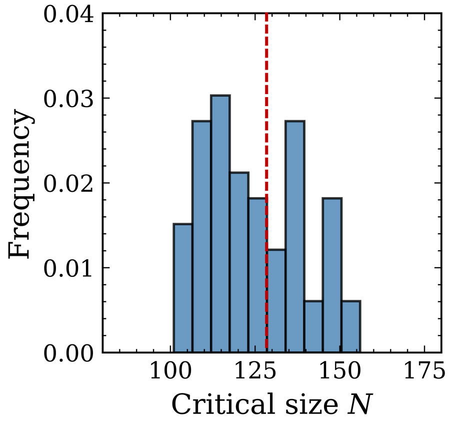


**Figure S5.** Critical cluster size distribution of agents (counted over 100 independent simulations), the red dashed line denotes the mean critical cluster size (N ≈ 128). The critical size is computed from the hysteretic profiles of non-induced phase separation, as shown in Figure S3.

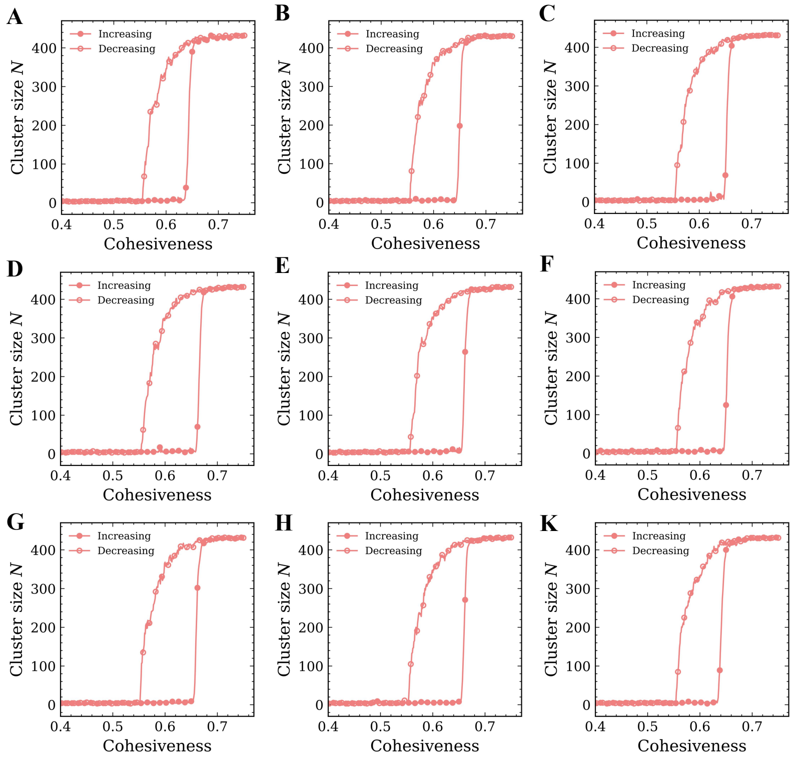


**Figure S6.** The cluster size of agents as a function of agent cohesiveness with the induction of CREs turned off in several independent runs.

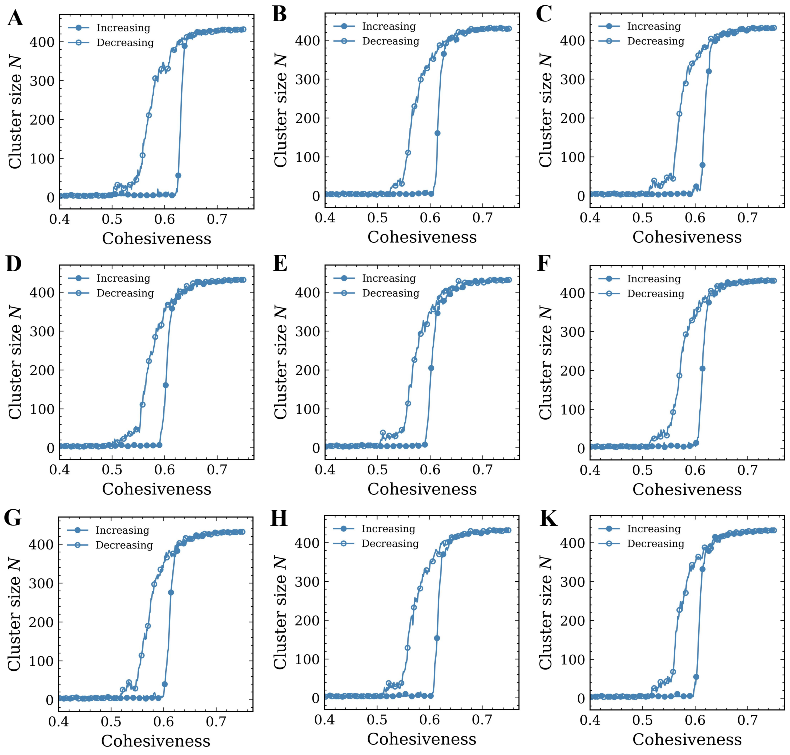


**Figure S7.** The cluster size of agents as a function of agent cohesiveness with the induction of CREs turned on in several independent runs.

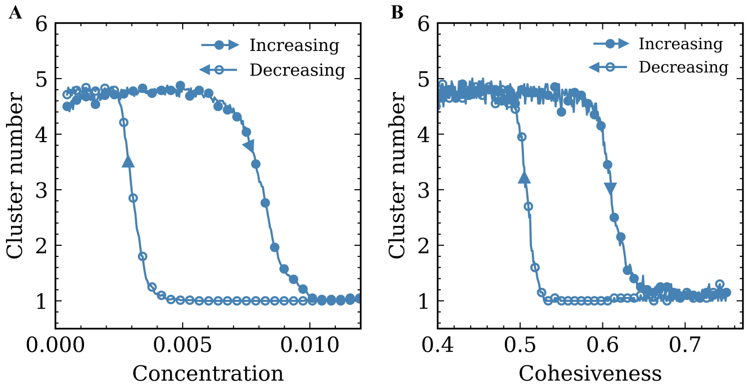


**Figure S8.** Clustering analysis of CREs in the (A) concentration- and (B) cohesiveness -dependent hysteretic simulations of phase separation (averaged over 100 independent simulations). Hysteretic behaviors have been observed. These results show the range of concentration or cohesiveness in which the CRE cluster would be stable once formed.

### 4. CRE clustering driven by induced LLPS

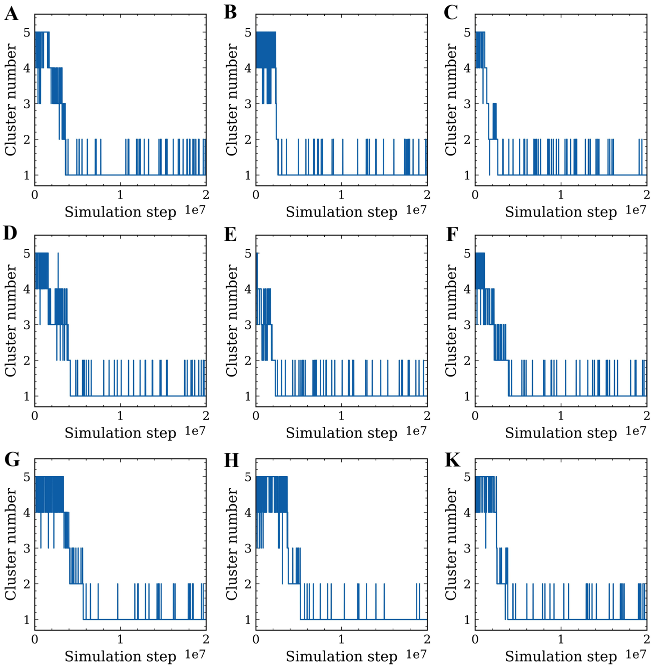


**Figure S9.** The cluster number of CREs as a function of simulation steps in several independent runs. The nonequilibrium simulations show stochasticity in the clustering dynamics.

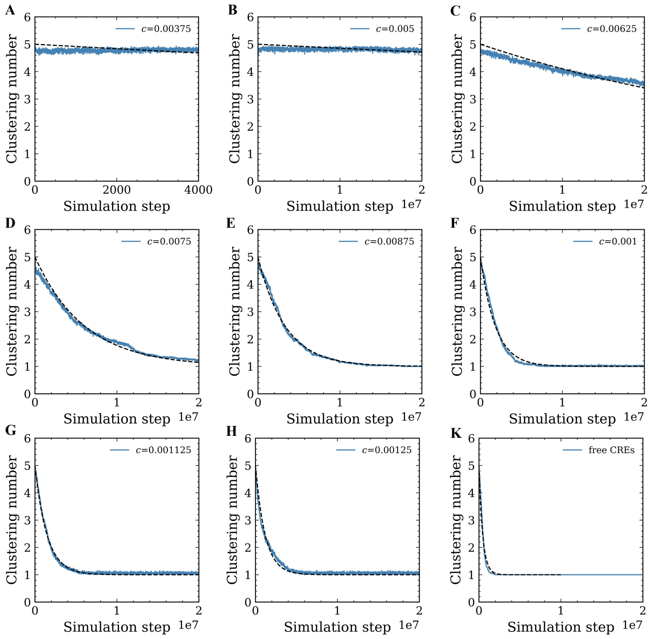


**Figure S10.** Exponential regressions of the CRE clustering profiles with different agent concentration. The simulation system of free CREs at $c = 0.01$ is used as a reference.

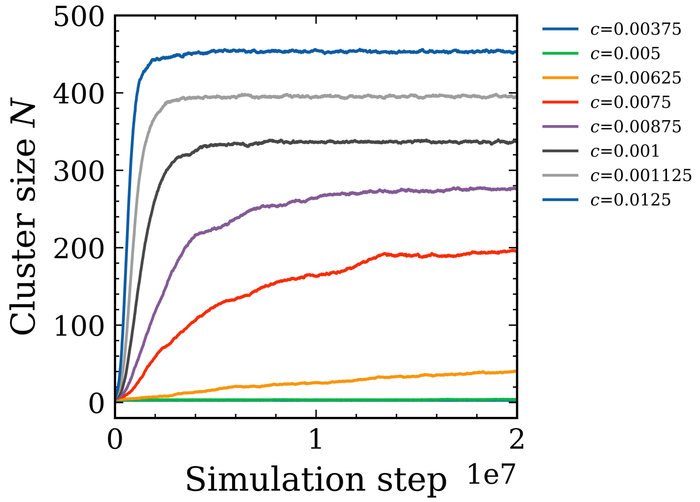


**Figure S11.** The largest cluster size of agents as a function of simulation step.

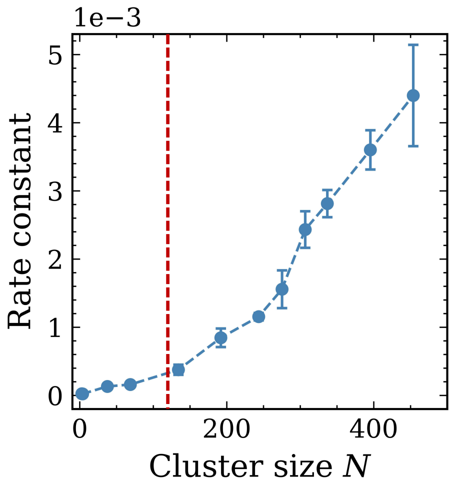


**Figure S12.** Correlation between the CRE clustering rate constant and the agent condensate size characterized by the agent number *N*. The red dashed line denotes the mean critical size, as depicted in Figure S4.

Figure S13 shows the whole process of CRE clustering with $\epsilon_{\text{CRE-agent}}$ = 0.74 kT. First, CREs are randomly distributed in the simulation box without contacts. After a long simulation period, two CREs bind together via agent bridges, this becomes a nucleation site to induce further phase separation. The agent cluster around the two CREs grows as more agents diffuse to the nucleation site over time, and CREs are recruited into the agent condensate through multivalent interactions.

To explore the effect of CRE-agent cohesiveness on CRE clustering dynamics, we increase the

$\epsilon_{\mathrm{CRE-agent}}$ to 0.85 kT and performed a control simulation, the whole process of CRE clustering is given in Figure S14. A different dynamical mechanism is found. Due to the stronger CRE-agent interaction, agents adsorb quickly on almost every CRE at first and form small clusters. The larger cluster then arises via the fusion of small clusters driven by the stochastic diffusion of proximal cluster, referred to as Brownian motion coalescence(*4*) (Figure S14B-C). Once the large cluster reaches to a certain size, the agents in other small clusters will "evaporate" and re-enter into the large cluster, known as Ostwald ripening(*5*) (Figure S10D-E). Finally, remaining CREs are recruited into the large cluster through multivalent interactions.

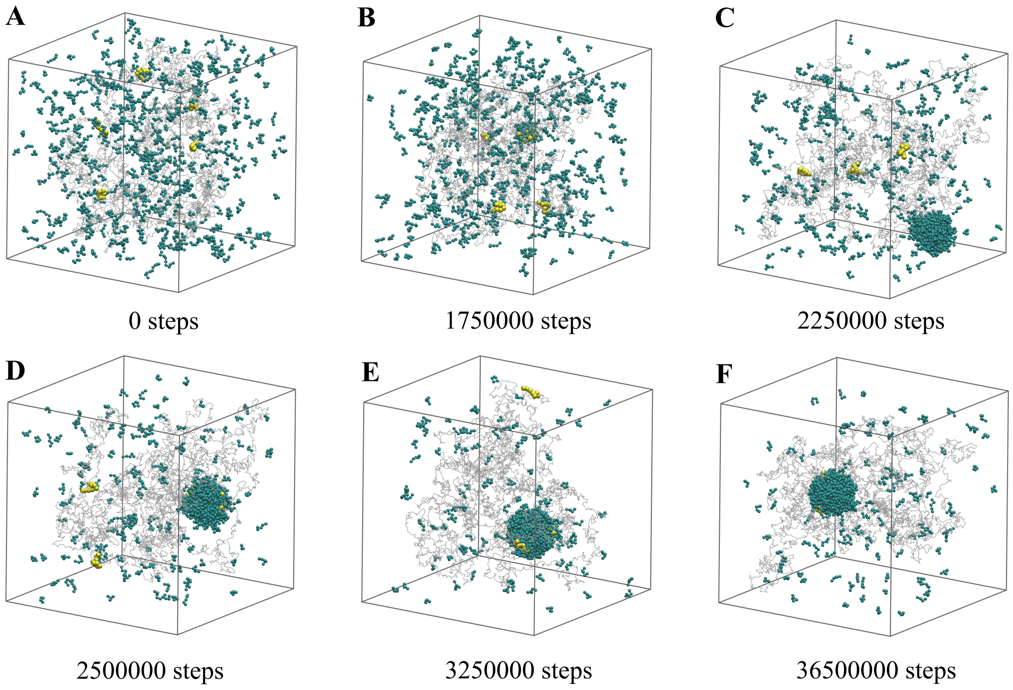


**Figure S13.** Time evolution of the CRE clustering driven by induced LLPS ($\epsilon_{\mathrm{CRE-agent}}$ = 0.74 kT), at the agent concentration $c$ = 0.01.

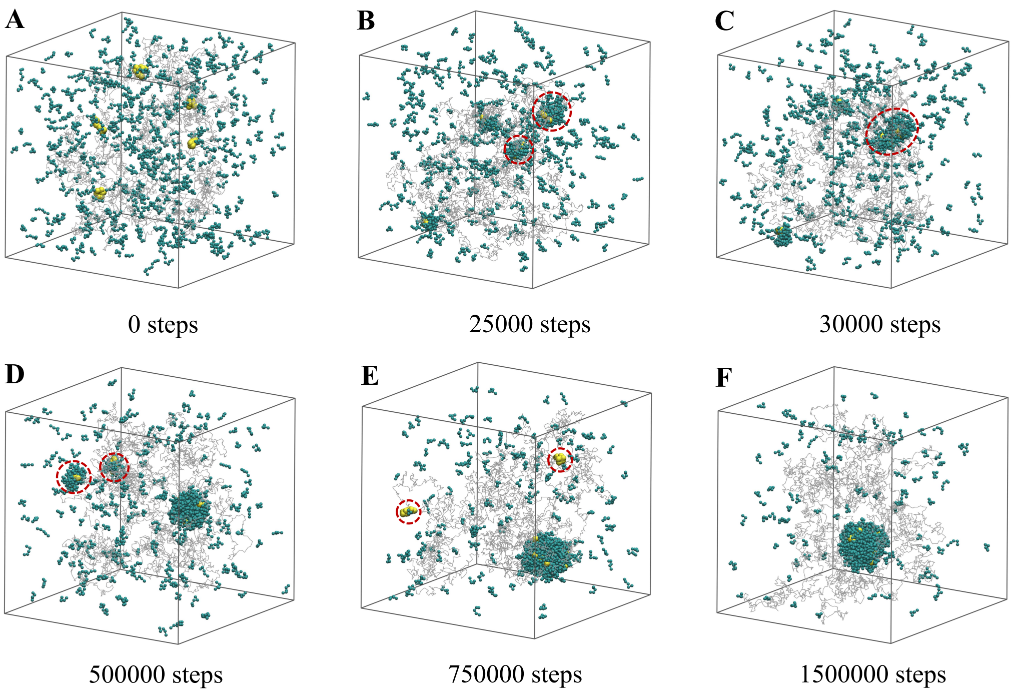


**Figure S14.** Time evolution of the CRE clustering driven by induced LLPS with stronger CRE-agent interactions ($\epsilon_{\mathrm{CRE-agent}}$ = 0.85 kT), at the agent concentration $c$ = 0.01.

### 5. CRE clustering driven by bridging effect

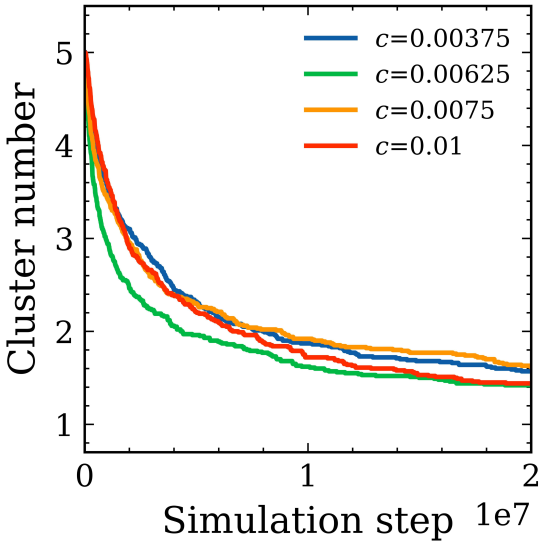


**Figure S15.** CRE clustering profiles (averaged over 100 independent simulations) of the bridging systems with different agent concentrations. The clustering rate is not proportional to the agent concentration and the clustering profile also does not follow exponent decay, which are much different from the cases of the induced LLPS systems.
